# A commensal adhesin enhances *E. coli* retention by mucin, while mucin desulfation by mucin-foraging bacteria enhances its transmigration through the mucus barrier

**DOI:** 10.1101/126672

**Authors:** Fitua Al-Saedi, Diana Pereira Vaz, Daniel H Stones, Anne Marie Krachler

**Affiliations:** University of Birmingham, School of Biosciences, Institute of Microbiology and Infection, Edgbaston, B15 2TT Birmingham, UK; Department of Microbiology and Molecular Genetics, University of Texas McGovern Medical School at Houston, Houston, TX, 77030, USA

**Keywords:** Multivalent Adhesion Molecule, mammalian cell entry (MCE) domain, adhesin, mucin, commensal, mucus-foraging, Bacteroides

## Abstract

Bacterial adhesion to host receptors is an early and essential step in bacterial colonization, and the nature of adhesion-receptor interactions determines bacterial localization and thus the outcome of these interactions. Here, we determine the host receptors for the Multivalent Adhesion Molecule from the gut commensal *E. coli* HS (MAM^HS^), which contains an array of seven mammalian cell entry (MCE) domains. The MAM^HS^ adhesin interacts with a range of host receptors, through recognition of a shared 3-O-sulfo-galactosyl moiety. This functional group is also found in mucin, a component of the intestinal mucus layer and thus one of the prime adherence targets for commensal *E. coli*. Mucin gels impede the motility of *E. coli* by acting as a physical barrier, and the barrier effect is enhanced by specific interactions between mucin and MAM^HS^ in a sulfation-dependent manner. Desulfation of mucin by pure sulfatase or the sulfatase-producing commensal *Bacteroides thetaiotaomicron* decreases binding of *E. coli* to mucin and increases attachment of bacteria to the epithelial surface, through interactions with surface-localized sulfated lipid and protein receptors. Together, our results demonstrate that the *E. coli* adhesin MAM^HS^ facilitates retention of a gut commensal by mucin, and suggest that the amount of sulfatase secreted by mucin-foraging bacteria inhabiting the same niche, such as *B. thetaiotaomicron*, may affect the capacity of the mucus barrier to retain commensal *E. coli*.

## INTRODUCTION

The gastrointestinal tract is covered by a mucus layer, which forms a semi-diffusive barrier protecting the underlying epithelium against microbial and chemical insults. The mucus layer consists of two parts, a cell-attached and a loose mucus layer (1), which is the habitat of the commensal flora (2). Both are mainly made up of the mucin Muc2, a large gel-forming glycoprotein that is modified with O-glycans, although levels and pattern of glycosylation vary with localization (3,4).

In healthy individuals, bacteria are prevented from accessing the inner mucus layer and epithelial surface, and are maintained at the luminal side of the mucus barrier (5). In contrast, the distal mucus layer, which undergoes fast renewal and turnover to maintain gut homeostasis (6) constitutes a distinct intestinal niche cohabited both by bacterial species that forage on mucus-derived glycans, such as *Bacteroides ssp.*, and species that lack mucolytic abilities including commensal *E. coli* (7,8). The composition and activity of the gut microbiota, as well as functional competition in this habitat can shape the mucus barrier in a way that determines microbe-host interactions (7-9). Aberrant production and O-glycosylation of Muc2 have been associated with increased bacterial penetration and thus elevated inflammatory responses both in rodent models and patients (10-12). The composition and localization of the gut microbiome is shifted in patients with inflammatory bowel diseases, and an overabundance of mucus-foraging species as well as a closer association of commensals including non-mucolytic *E. coli* has been reported (13). Although a causality between these two observations has not been firmly established, it has been hypothesized that the enhanced production of mucinolytic enzymes leads to compromised barrier integrity, which facilitates the relocation of microbes in close proximity of the epithelium (14). However, the molecular mechanisms underpinning the interplay between microbial-mucus interactions and bacterial localization in the intestinal habitat are subject to ongoing investigation.

Here, we set out to study the host receptors of an adhesin belonging to the family of Multivalent Adhesion Molecules (MAMs) from the gut commensal *E. coli* strain HS. MAMs are abundant in Gram-negative bacteria, and in the context of pathogens have been shown to initiate adhesion to host tissues and infection (15,16). The *Vibrio parahaemolyticus* MAM (MAM7) has been shown to bind phosphatidic acids, anionic lipids found at the host membrane, and use the extracellular matrix protein fibronectin as a co-receptor (17). It was further shown that MAM7-binding to host cells competitively precludes other bacteria from adhering (15,18), and that this could be utilized as a strategy to combat infection *in vivo* (19). Our recent work showed that binding of the MAM homolog from commensal *E. coli* HS (or short, MAM^HS^), could also competitively inhibit bacterial attachment to host cells (20). However, whether it targets the same host receptors as MAM7 was unclear and its functionality could not be deducted based on sequence, due to low sequence conservation.

Here, we show that the MAM^HS^ adhesin derived from the commensal *E. coli* strain HS binds to anionic host lipids, but in contrast to *V. parahaemolyticus* MAM7 has a preference for sulfated lipids. MAM^HS^ also interacts with a range of host protein receptors, through a conserved 3-O-sulfo-galactosyl moiety that is conserved between sulfoglycosylated proteins and sulfatide (3-O-sulfogalactosylceramide). The interaction of MAM^HS^ with sulfated mucin competitively inhibits attachment of *E. coli* to epithelial cells, and enhances the retention of *E. coli* by a mucin gel. This barrier effect is significantly decreased by mucin desulfation, such as by the mucin forager *Bacteroides thetaiotaomicron*.

Our results suggest that the physical barrier effect of mucus can be modulated by specific interactions between commensal adhesins and mucin, and that the interplay between mucin and microbes cohabiting the same intestinal niche as *E. coli* may modulate the localization of commensals within the gut.

## RESULTS

### Multivalent Adhesion Molecules from different bacteria show conserved binding to host lipids but show preference for different anionic head groups

We have previously shown that the Multivalent Adhesion Molecule (MAM) from the commensal *E. coli* strain HS (MAM^HS^) competitively inhibits pathogen adherence to human cells (20). However, the protein sequence of *E. coli* MAM^HS^ is quite different to the sequence of *Vibrio parahaemolyticus* MAM7, the first characterized MAM which uses the host lipid phosphatidic acid and the extracellular matrix protein fibronectin as receptors (15,17). To determine ligand binding specificity, we first tested the lipid binding properties of MAM^HS^ using lipid overlay assays. While MAM7 from *V. parahaemolyticus* bound to phosphatidic acid (15), *E. coli* MAM^HS^ bound to sulfatide (Fig. 1A). Like phosphatidic acid, sulfatide (3-O-sulfogalactosylceramie) comprises of a negatively charged head group which is necessary for binding. However, no binding was detected to ceramide, which is structurally identical to sulfatide but lacks the sulfo-galactosyl moiety (Fig. 1B). To quantitate binding of both MAM7 and MAM^HS^ to lipids, we used a modified indirect quantitative ELISA and chemoluminescence detection. Wells of a high-bind microtiter plate were coated with sulfatide, ceramide, phosphatidic acid or were left uncoated. GST-MAM7 or GST-MAM^HS^ were added at indicated concentrations and binding quantified by incubation of plates with GST-antibody followed by incubation with a secondary HRP-coupled antibody and chemoluminescence measurements. While MAM7 showed strong binding to phosphatidic acid, it only bound weakly to sulfatide and showed no significant binding to ceramide or uncoated wells (Fig. 1C). In contrast, MAM^HS^ bound to sulfatide with high affinity, but displayed much lower binding to phosphatidic acid and ceramide (Fig. 1D). These data demonstrate that while both MAMs specifically bind to anionic host lipids, *V. parahaemolyticus* MAM7 preferentially binds to phosphatidic acid, which features a glycerophosphate head group, while *E. coli* MAM^HS^ prefers sulfatide, featuring a galactosyl-sulfate moiety, as receptor.

**Fig. 1.**
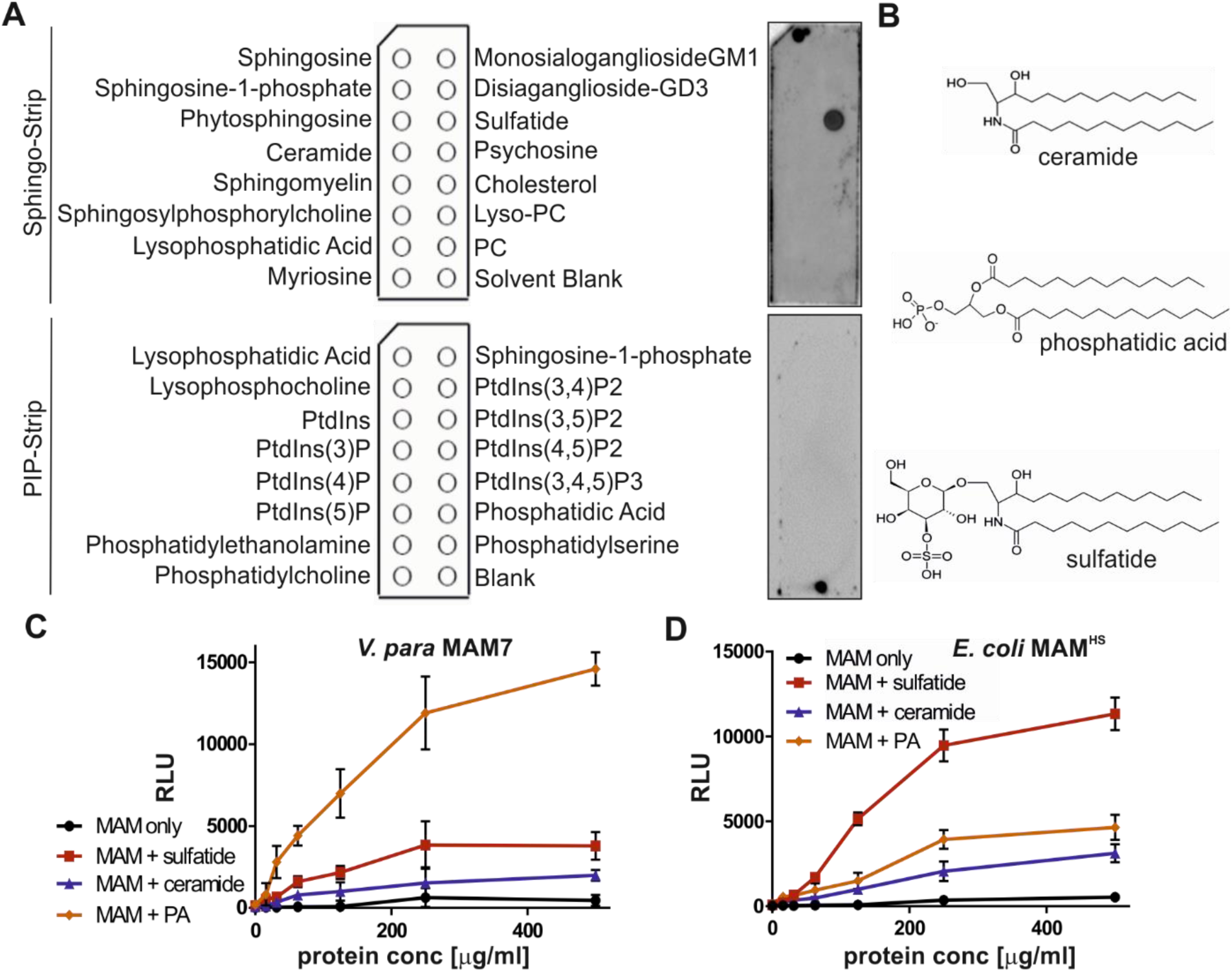
*V. parahaemolyticus* MAM7 and *E. coli* MAM^HS^ show different binding preferences for host lipid receptors. (A) The lipid binding profile of *E. coli* GST-MAM^HS^ was determined by lipid overlay assays. 10 µM recombinant pure protein were incubated with Sphingo-Strips or PIP-Strips, and bound proteins detected by probing membranes with α-GST and α-mouse HRP antibodies, and developing with ECL detection reagent. (B) Comparison of chemical structures of ceramide, phosphatidic acid, and sulfatide head groups. Interactions between *V. parahaemolyticus* MAM7 (C) or *E. coli* MAM^HS^ (D) and lipids were quantified using plate assays. Lipids were immobilized in wells and bound protein was quantified by probing wells with a-GST and a-mouse HRP antibodies, and developing with ECL detection reagent. Results shown are means ± SEM from three independent experiments.

### The commensal adhesin MAM^HS^ from E. coli binds to sulfated host proteins and lipids through specific recognition of a shared 3-o-sulfo-galactosyl moiety

In parallel to testing its lipid binding profile, we also conducted pull-down experiments with epithelial cell lysates to determine if *E. coli* MAM^HS^ would recognize host proteins as adhesion receptors. GST-MAM^HS^ was used as a bait to pull down bound proteins from Hela cell lysate, and GST alone was used as a negative control. SDS-PAGE revealed the presence of seven specific bands, corresponding to MAM, GST, and five putative interacting proteins (Fig. 2). Bound host proteins (corresponding to bands 2-6) were identified as perlecan, mucin, fibronectin, collagen IV and laminin as top hits. With the exception of mucin which coats mucosal surfaces, all of these proteins are part of the extracellular matrix and all can either be glycosulfated, or in the case of laminin, bind directly to sulfatides (21,22). This led us to test the binding specificity of MAM^HS^ towards glycosulfates.

**Fig. 2.**
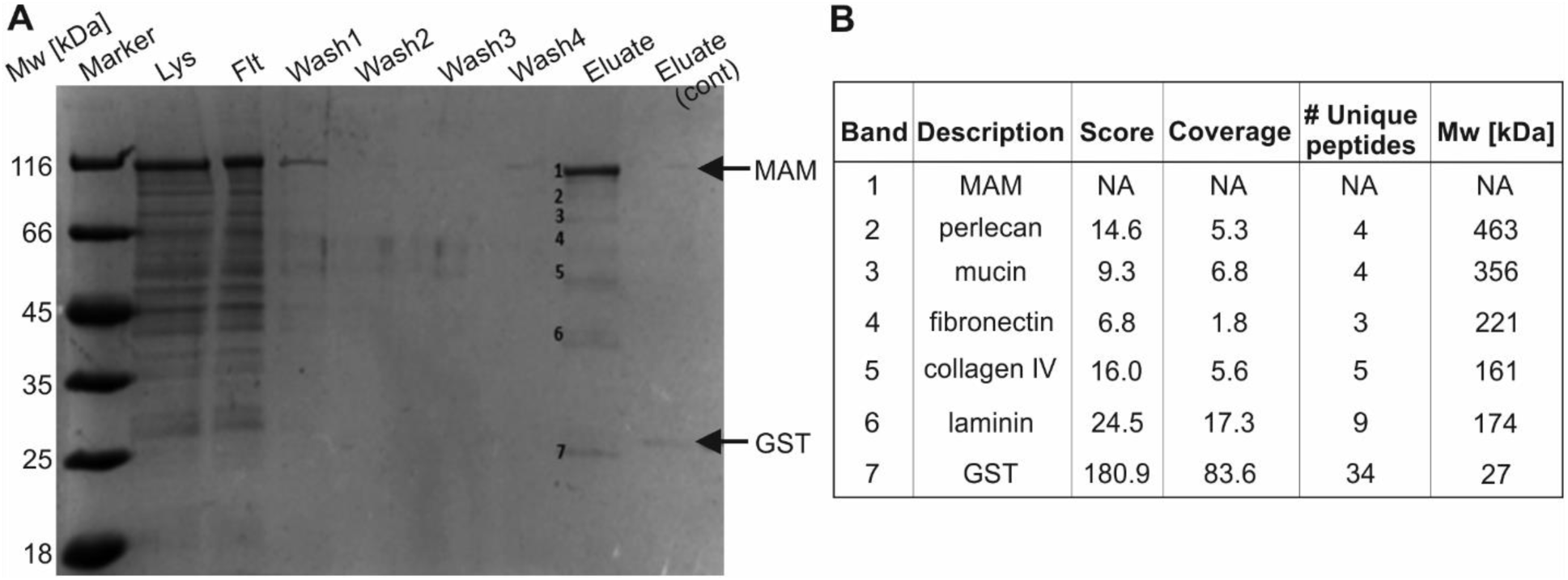
*E. coli* MAM^HS^ interacts with multiple sulfated host proteins. (A) Pull-down of Hela lysate with purified *E. coli* GST-MAM^HS^. Lysate (Lys), flowthrough (Flt), wash and eluate fractions were resolved by SDS-PAGE. A control experiment was done using GST only and eluate from this experiment was run on the same gel (eluate cont.). Molecular weights of GST-MAM^HS^ and GST control are indicated by arrows. Eluted bands 1-7 were cut out and subjected to tryptic digest and protein ID by LC-MS/MS. (B) Top hits for protein bands 1-7 depicted in (A).

Competitive attachment assays testing the attachment of BL21 expressing MAM^HS^ either in the absence or presence of small molecule inhibitors demonstrated MAM-mediated adhesion was specifically abolished by lactose-3-sulfate. Adhesion was unaffected by the presence of µM concentrations of lactose, galactose, galactose-6- sulfate, N-acetyl-glucosamine, or N-acetyl-glucosamine-6-sulfate (Fig. 3). Lactose-3-sulfate shares the 3-O-sulfo-galactosyl moiety found in sulfatide. These results show that recognition is glycol-specific, but also specific to the position of the sulfoglycosylation, since galactose-6-sulfate did not inhibit binding.

**Figure 3.**
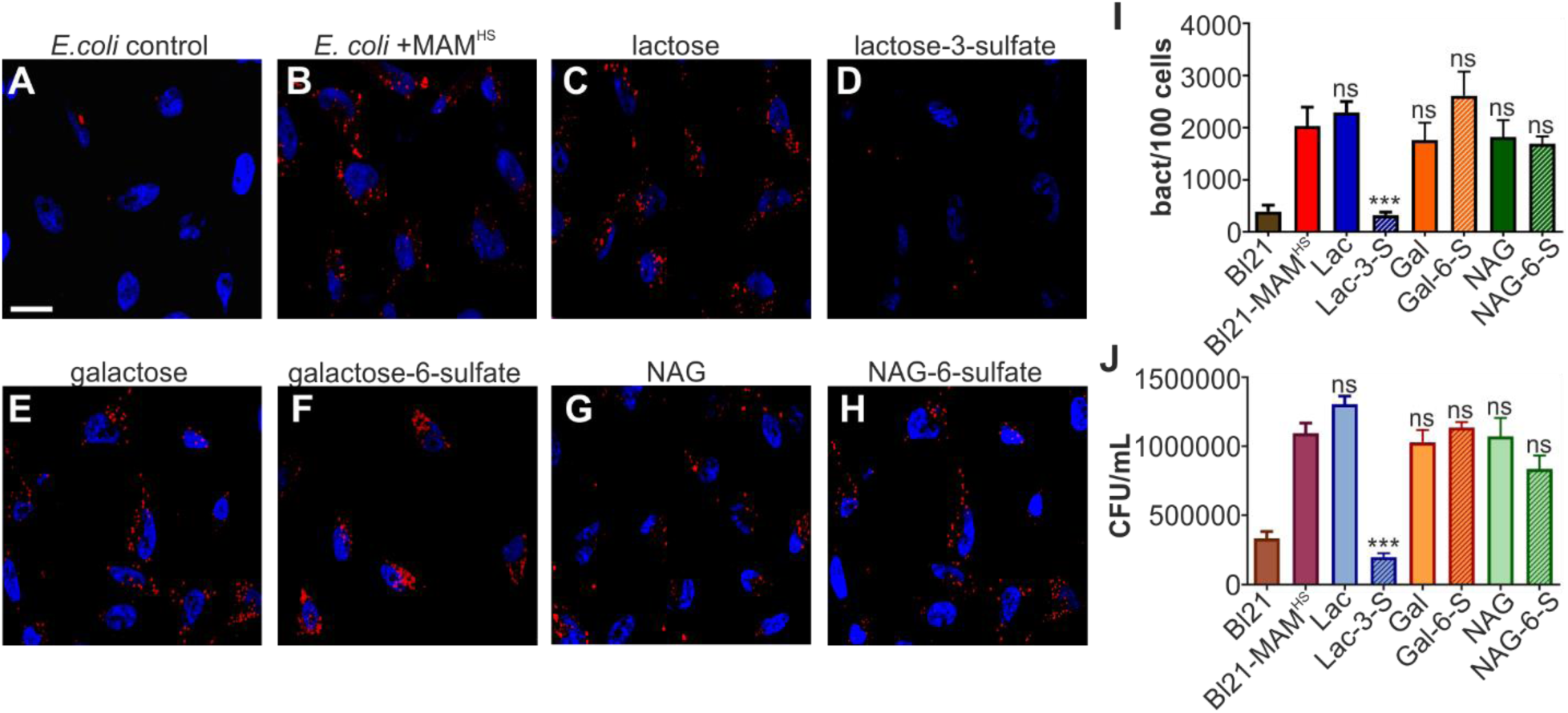
A 3-O-galactosyl moiety is the common functional group recognized on host receptors by *E. coli* MAM^HS^. Bacterial attachment of mCherry-expressing BL21 control (A) or BL21 expressing mCherry and MAM^HS^ (B) to Hela epithelial cells. Bacterial attachment of BL21 expressing mCherry and MAM^HS^ in the presence of 200 µM lactose (C), lactose-3-sulfate (D), galactose (E), galactose-6-sulfate (F), N-acetylglucosamine (G) or N-acetylglucosamine-6-sulfate (H). Cells were fixed and stained with Hoechst (blue) and bacteria are visible in red. Scale bar, 10 µm. Bacterial attachment was quantified by image analysis (I) and values plotted are means ± sem from at least 100 cells per condition (n=3). Bacterial attachment was quantified by serial dilution plating of Triton-lysed cells (J) and values represent means ± sem (n=3). Significance was determined using student’s two-tailed t-test. (***) indicates p≤0.001, ns indicates p≥0.05.

### Sulfation-dependent binding to mucin prevents MAM^HS^-mediated adherence to epithelial cells

Of all host proteins we identified, mucin is one of the earliest targets encountered by bacteria entering the gastrointestinal tract and hence constitutes a prime target for bacterial adherence. Mucin is a major component of the mucosal layer of the gastrointestinal tract, while all other identified targets are either localized at the epithelial surface or basement membrane. Hence, we focused our further studies on the interaction between MAM^HS^ and mucin.

To confirm if MAM^HS^ and mucin directly interact, we coated microtiter plates with either mucin from porcine stomach type II, type III, or mucin type I-S from bovine submaxillary glands, and quantitated binding of GST-MAM^HS^ using a modified indirect ELISA and chemoluminescence detection (Fig. 4). GST-MAM^HS^ strongly bound both mucin type II and III from porcine stomach (Fig. 4A,B), while no binding was detected to mucin from bovine submaxillary glands (Fig. 4C). All three mucins contain the same Muc2 core structure, but differ in their level and quality of glycosylation, with mucin type I-S carrying shorter O-linked glycans and being most heavily sialylated. These data suggest that MAM^HS^ specifically recognizes glycosylation patterns on mucin.

**Figure 4.**
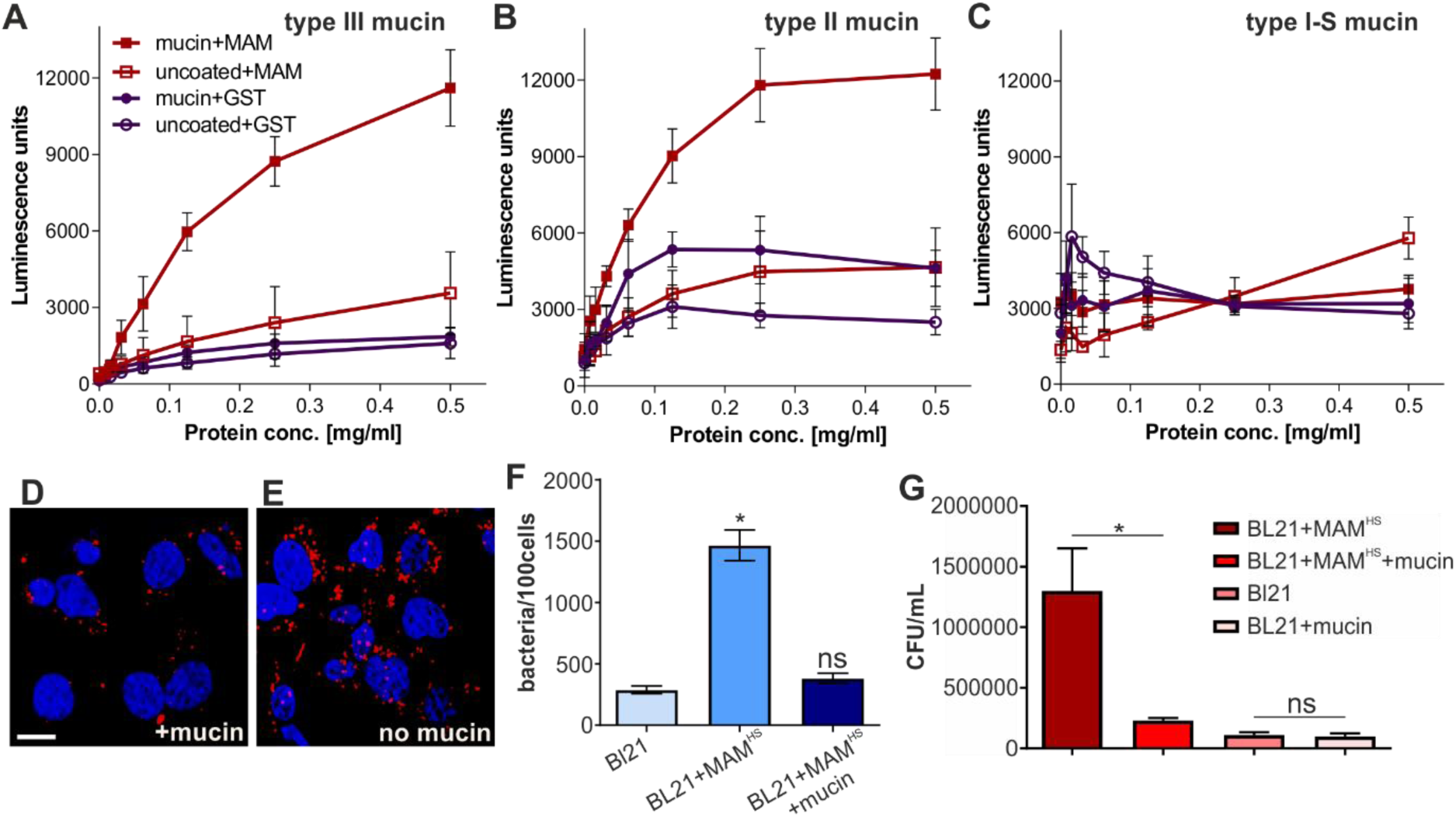
*E. coli* MAM^HS^ - mucin interaction competes for cell surface binding. Binding of purified GST-MAM^HS^ or GST (control) to mucin from porcine stomach type III (A), mucin from porcine stomach type II (B) and mucin type I-S from bovine submaxillary glands (C) was quantified using a-GST and a-mouse HRP antibodies coupled with chemoluminescence detection. Wells contained either 5 µg mucin or were left uncoated. GST-MAM or GST protein were added at concentrations between 0-0.5 mg/ml. Values represent means ± sem (n=3). Attachment of BL21-MAM^HS^ + mCherry to Hela cells with (D) or without (E) prior incubation of bacteria with mucin. Blue – Hoechst, red – mCherry *E. coli*. Scale bar, 10 µm. (F) Quantification of attached bacteria from experiment shown in D and E. Values are means ± sem from at least 100 cells per condition (n=3). Significance was determined using student’s two-tailed t-test. (*) indicates p≤0.05, ns indicates p≥0.05. (G) Bacterial attachment was quantified by serial dilution plating of Triton-X100 lysed cells. Values represent means ± sem (n=3). Significance was determined using student’s two-tailed t-test. (*) indicates p≤0.05, ns indicates p≥0.05.

Next, we tested if the interaction between *E. coli* expressing MAM^HS^ and mucin would affect the bacteria’s ability to adhere to epithelial cells. Pre-incubation of BL21 MAM^HS^ with mucin type II decreased bacterial attachment to Hela cells to background levels (BL21 without MAM^HS^), while the non-specific adherence of BL21 without MAM^HS^ was not affected by incubation of bacteria with mucin (Fig. 4D-G). These data demonstrate that interaction of MAM^HS^ with mucin impedes bacterial attachment to the epithelial surface, and this effect is not due to physical retention of bacteria by mucin.

Mucin, like the other protein receptors for MAM^HS^ we identified, is glycosylated and glycosulfation constitutes one of the main glycan modifications found on gastrointestinal mucin. Sulfated residues on intestinal mucin chiefly include N-acetylglucosamine, N-acetyl-galactosamine, and galactose (23-25). Since our data showed that galactose-3-sulfate is an inhibitor of bacterial attachment (Fig. 4D), and a common moiety found both on sulfatides and sulfoglycosylated proteins such as mucin, we tested if mucin sulfation was required for the interaction between MAM^HS^ and mucin. Treatment of mucin type II with purified sulfatase from *Helix promatia* released sulfate in a concentration-dependent manner (Fig. 5A). Desulfated mucin was then used to test for GST-MAM^HS^ binding by a modified indirect ELISA, and desulfation decreased the binding of GST-MAM^HS^ in a dose-dependent manner (Fig. 5B). Desulfated mucin also lost the ability to prevent *E. coli* expressing MAM^HS^ from attaching to the epithelial surface, as determined both by image analysis (Fig. 5C-E) and dilution plating of bacteria attached to host cells in the presence and absence of desulfated mucin (Fig. 5F).

**Figure 5.**
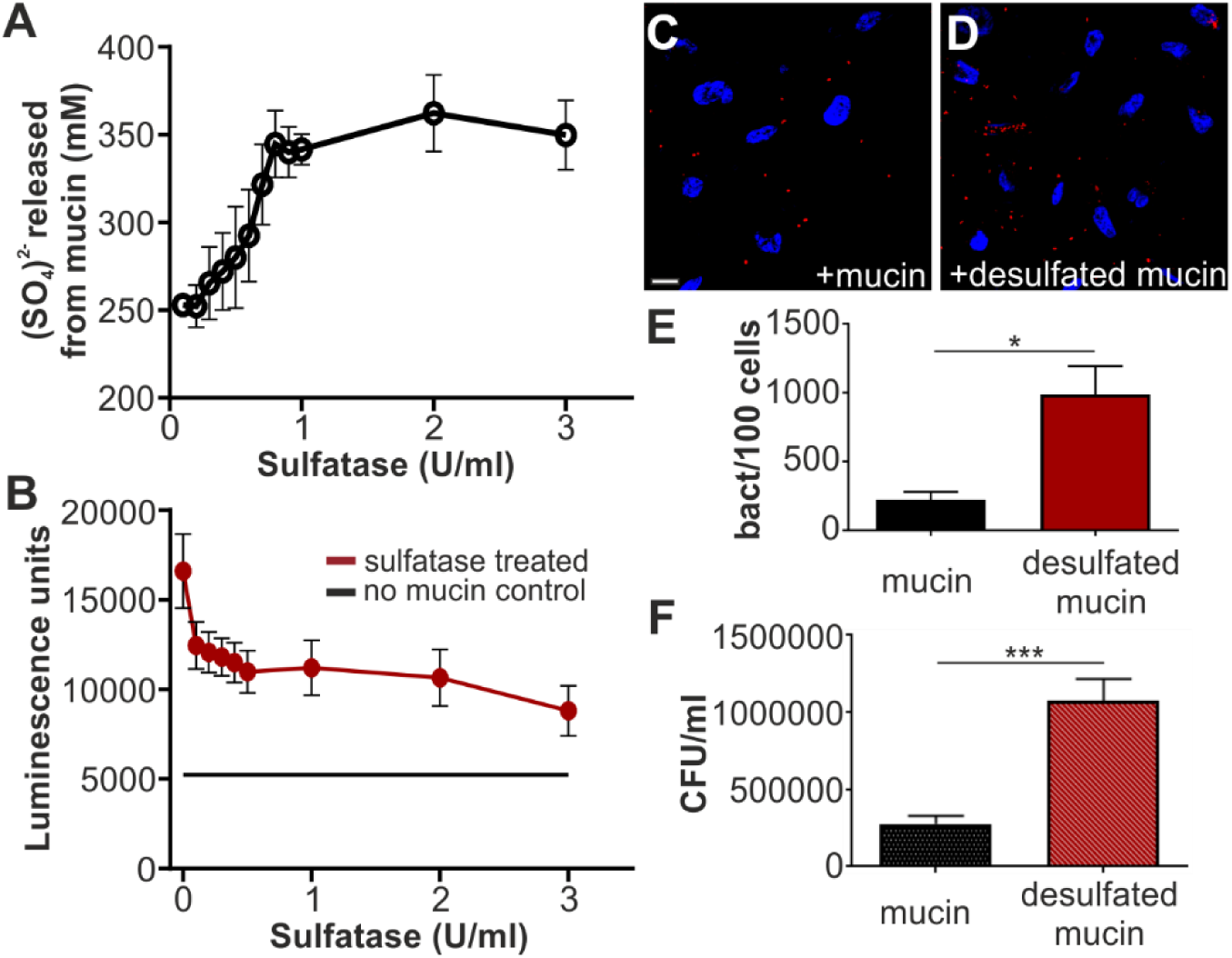
*E. coli* MAM^HS^ binding to mucin is sulfation-dependent. (A) Treatment with sulfatase from *Helix promatia* releases sulfate from mucin in a concentration-dependent manner. (B) Desulfation of mucin with 100 µl sulfatase from *Helix promatia* (activities as indicated) decreases GST-MAM^HS^ binding to mucin from porcine stomach type II. Bacterial attachment BL21 expressing mCherry and MAM^HS^ to Hela epithelial cells in the presence of 50 µg/ml type II mucin (C) or desulfated mucin (D). Cells were fixed and stained with Hoechst (blue) and bacteria are visible in red. Scale bar, 10 µm. Bacterial attachment was quantified by image analysis (E) and values plotted are means ± sem from at least 100 cells per condition (n=3). Bacterial attachment was quantified by serial dilution plating of Triton-lysed cells (F) and values represent means ± sem (n=3). Significance was determined using student’s two-tailed t-test. (*) indicates p≤0.05, (***) indicates p≤0.001.

While *E. coli* is unable to desulfate mucin, *Bacteroides* species, which inhabit the same gastrointestinal niche as *E. coli*, are prolific secretors of sulfatases. We therefore asked whether treatment of mucin with the secretions of the human gut symbiont *Bacteroides thetaiotaomicron* would alter MAM^HS^ binding to mucin. We determined sulfatase activity in cell-free supernatants isolated from *B. theta* cultured in BHI for 48 hours, as well as a sulfatase-deficient mutant (*B. theta* anSME, Fig. 6A). We then treated type II mucin with 100 µl *B. theta* supernatant, corresponding to a sulfatase activity of 0.1 U, which we had previously determined to be a sufficient activity to cause loss of MAM^HS^ binding to mucin (Fig. 5B). Treatment of mucin with supernatant from wild type *B. theta*, caused a decrease of MAM^HS^ binding to mucin, which increased with increasing exposure time of mucin to the supernatant prior to incubation with MAM^HS^ and eventually lead to a complete loss of binding (Fig 6B). Treatment of mucin with supernatant from the sulfatase-deficient *B. theta* anSME mutant caused a significant, albeit much smaller loss in MAM^HS^ binding after extended incubation of supernatant with mucin. This can be attributed to B*. theta*’s ability to degrade mucin during extended exposure. Taken together, these experiments show that binding of MAM^HS^ to mucin depends on the presence of sulfoglycan modifications on mucin, and that the exposure of mucin to sulfatase-secreting *B. theta* decreases sulfation and thus the ability of mucin to bind MAM^HS^, which enables bacteria to attach to the epithelial surface.

**Fig. 6.**
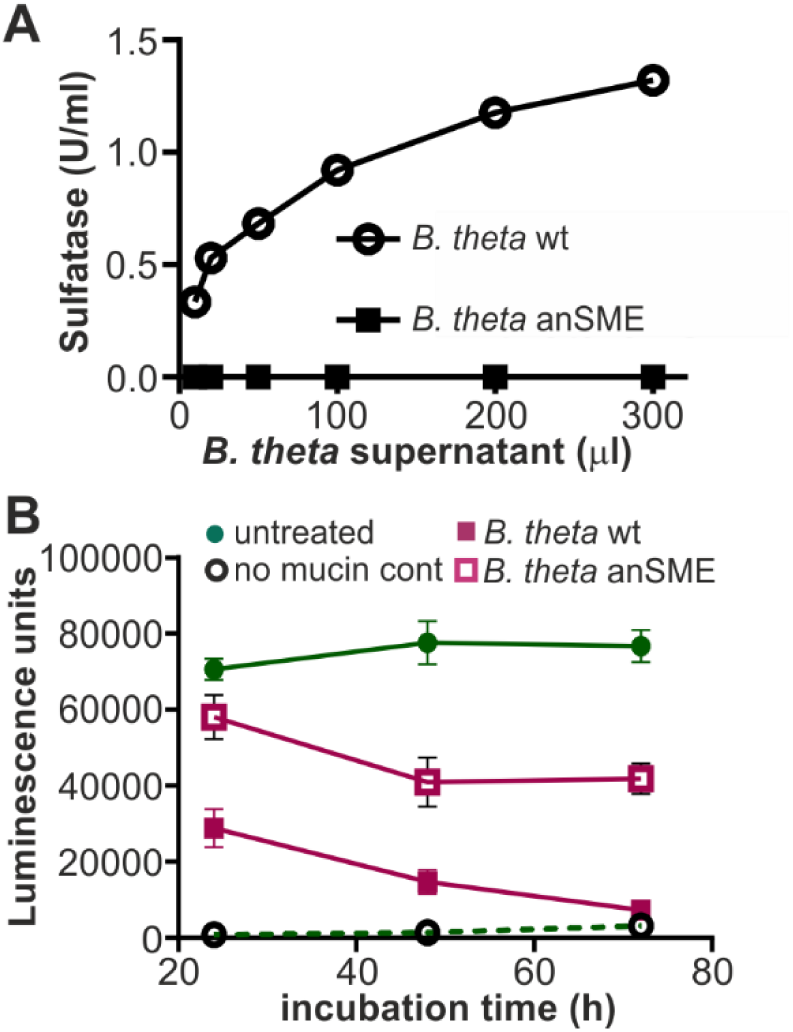
*E. coli* MAM^HS^ binding to mucin is decreased by exposure of mucin to sulfatase-secreting *B. thetaiotaomicron*. (A) Sulfatase activity of supernatants from *B. theta* wt (circle) and anSME mutant (square) were quantified using p-nitrocatechol sulfate as substrate. (B) Mucin type II from porcine stomach was treated with 100 µl (1U/ml) of supernatant from *B. theta* wt (pink squares) or anSME mutant (empty squares) for the indicated times, and used in a plate assay to quantify GST-MAM^HS^ binding. As control, GST-MAM^HS^ binding to untreated mucin (green circles), or binding to uncoated wells (empty circles) was measured. All values plotted are means ± sem (n=3).

### Mucin desulfation helps E. coli to transmigrate a mucin gel simulating the mucus barrier

The above-described experiments suggest that the mucin-desulfating activity of sulfatase-producing bacteria cohabiting the same niche as *E. coli* affect the ability of *E. coli* to bind to and be retained by intestinal mucus. Although these experiments conclusively demonstrate competition for *E. coli* MAM-dependent binding to mucin or the epithelial surface, they use soluble mucin and thus do not adequately reflect the situation in the GI tract, where a continuous mucus layer precludes access of bacteria to the underlying epithelium. In such a situation, the ability of bacteria to reach the epithelium depends on both specific binding to mucin, as well as the impact of mucin-modification on structural integrity of the mucin layer. We attempted to simulate this environment by using long-term cultured mucin-producing colonic epithelial cells (HT29-MTX). However, despite our attempts to optimize culture conditions, including mucin supplementation and simulated flow, we only observed a patchy distribution of mucus and not a continuous mucus layer. Thus, we sought to simulate this more complex scenario by reconstituting a mucin gel to simulate the barrier effect of the intestinal mucus layer, and test how sulfation would impact *E. coli* transmigration through this barrier.

*E. coli* rapidly migrated through uncoated transwells (pore size 3 µm) and motility was unaffected by the presence or absence of MAM^HS^ (Fig. 7A). Coating of transwells with a mucin gel impeded the transmigration of *E. coli*, but retained *E. coli* expressing MAM^HS^ more efficiently, suggesting the mucin layer constitutes a physical barrier but its effect is enhanced through specific interactions between mucin and the MAM^HS^ adhesin. Treatment of mucin with *Helix promatia* sulfatase increased the transmigration of *E. coli* expressing MAM^HS^ to levels seen with BL21 control bacteria, while mucin desulfation had no significant effect on the transmigration of *E. coli* without MAM^HS^ (Fig. 7A). This suggests that desulfation decreases the retention of bacteria expressing the commensal adhesin MAM^HS^, while it has no discernible impact on the mechanical barrier effect of mucin gels. These results were recapitulated when mucin was treated with *B. theta* supernatant, although in this case the permeability change is the result of both desulfation and partial mucin degradation by *B. theta* (Fig. 7B).

**Fig. 7.**
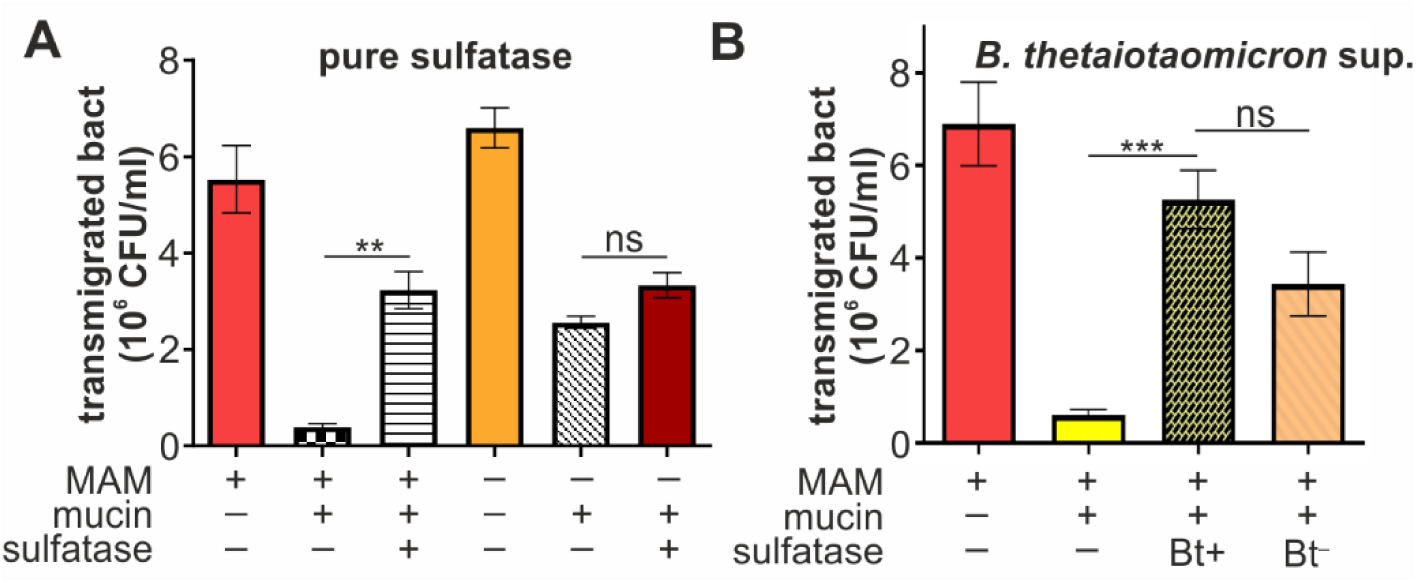
Mucin desulfation helps *E. coli* to transmigrate a mucin gel simulating the mucus barrier. (A) BL21 expressing MAM^HS^ (MAM +) or carrying empty vector (MAM -) were added to the top compartment of a transwell either coated with type II mucin (mucin +) or left uncoated (mucin -). Mucin used for coating was either desulfated with sulfatase from *Helix promatia* (sulfatase +) or left untreated (sulfatase -) prior to coating transwells. Bacterial transmigration (recovery from bottom well following 2 hours of incubation at 37 °C) was measured by serial dilution and plating. (B) Experiments were conducted as described in (A), but mucin was either treated with supernatant from wild type *B. theta* (Bt+) or from the sulfatase-deficient *B. theta* anSME mutant (Bt-) prior to coating transwells. Values represent means ± sem (n=3). Significance was determined using student’s two-tailed t-test. (***) indicates p≤0.001, (**) p≤0.01, ns indicates p≥0.05.

## DISCUSSION

The protective role of the intestinal mucus layer as a barrier against microbial invasion of the underlying epithelium, and against inflammatory responses resulting from bacteria-epithelial interactions, is well recognized (26,27). Similarly, it has been shown that the thickness of the mucus layer, and the integrity and glycosylation of its main component, the mucin protein Muc2, are key to maintaining the barrier effect (10-12). Although increased microbial penetration of the mucus layer has been linked to a loss in mucin glycosylation, the mechanisms behind these changes are not well understood.

Here, we report the adhesin MAM^HS^ found in commensal *E. coli*, interacts with mucin in a sulfation-dependent manner (Fig. 5). Through a 3- O-sulfo-galactosyl moiety shared between sulfated mucin and other host-derived proteins (Fig. 2-3), as well as host sulfatides (Fig. 1), MAM^HS^ can attach either to mucin or the host epithelium (Fig. 4).

Pull-down experiments with purified GST-MAM^HS^ as a bait identified five host epithelial proteins specifically interacting with the adhesin. All identified binding partners are extracellular proteins, and have been reported to be heavily glycosylated and sulfoglycosylated (28-30), or to bind to sulfated ligands, such as in the case of laminin, which binds to sulfatides (21,22). Further analysis of competitive inhibition with sugars or sulfosaccharides showed that the common moiety recognized by MAM^HS^ was galactose-3-sulfate, which is also found in sulfatide.

Sulfatide is an abundant host cell lipid, accounting for 4 % of total membrane lipid content (31,32). Sulfatides participate in a wide range of cellular functions, including protein trafficking, cell adhesion and aggregation, as well as immune responses (33). Additionally, both viruses and bacteria have been found to utilize sulfatides as host receptors for adherence, including enterotoxigenic *E. coli* (34). However, the identification of 3-O-sulfo-galactosyl as a common binding moiety promoting bacterial attachment to both host lipids and proteins, to our knowledge, has not been reported previously.

Mucin is a commonly described mediator of bacterial adherence, which is unsurprising given its abundance in the gastrointestinal tract and on other mucosal surfaces throughout the body, and its role in maintaining the local microflora. However, mucin displays a large variety of glycoepitopes, and as such bacteria have evolved a diverse range of adhesin-mucin interactions. Specific interactions have been described between *Pseudomonas aeruginosa* and Lewis x, sialyl-Le(x) and sulfosialyl-Le(x) glycoepitopes, and between *Helicobacter pylori* and Lewis (b) on respiratory and gastric mucins, respectively (35,36). Targeting of human blood type-A, B, or H-antigens modifying intestinal mucin by *Lactobacillus* has also been reported (37,38). *Lactobacillus reuteri* has been shown to recognize mucin in a sulfation-dependent manner, although the exact functional epitope remains elusive (39). The interactions between microbes and mucin oligosaccharides have been shown to facilitate bacterial clearance and to inhibit colonization of epithelial cells (40).

Sulfation of colonic mucins has been shown to play a protective role in animal models of colitis (23), and reduced sulfation has been associated with impaired barrier function and with infectious disease (41,42). Our data show that the sulfation status of mucin also directly impacts bacterial adherence to, and thus retention of bacteria by the mucin gel. Sulfatases produced by anaerobes colonizing the intestinal tract, such as the commensal *Bacteroides thetaiotaomicron*, are essential for mucosal foraging, and have a direct impact on the chemical composition and architecture of the mucus layer (43,44). Enhanced sulfatase production by the intestinal microbiota has been detected in fecal samples from patients suffering ulcerative colitis (45).

Our data suggest that the level of sulfatase production, and thus mucin desulfation, can act to modulate the adherence of commensals to mucin, and directly impact their retention by the mucus barrier. While the experiments herein focused on the biochemical determinants of MAM^HS^-receptor interactions, future work will aim to directly assess the impact of sulfatase-production on commensal adherence in a physiologically relevant model. Such experiment will determine if the observed changes in adherence in response to desulfation described herein, may have an impact on bacterial localization and mucosal inflammation *in vivo*.

## EXPERIMENTAL PROCEDURES

### Bacterial strains and growth conditions

Bacterial strains used in this study were *Escherichia coli* HS (accession ABV06236.1;GI:157066981) and *Bacteroides thetaiotaomicron* (wild type strain VPI-5482) and a derivative strain lacking the anaerobic sulfatase-maturating enzyme (anSME), ΔBT0238, that is unable to produce functional sulfatases (43). *E. coli* BL21 expressing MAM^HS^ or *E. coli* BL21 carrying empty pBAD (control) were grown in LB medium at 37 °C. *B. thetaiotaomicron* was grown in Brain Heart infusion (BHI) medium at 37 °C in an anaerobic jar.

### Protein expression and purification

10 ml of LB containing 100 μg/ml ampicillin was inoculated with a colony of *E. coli* BL21 containing the recombinant plasmid (p GEX-4T3 – MAM7ΔTM or MAM^HS^ΔTM) for production of GST-MAM7 or GST-MAM^HS^ protein, or *E. coli* BL21 containing pGEX-4T3 plasmid for production of GST protein, and incubated at 37 °C for 16 hours with shaking. Protein expression and purification was then carried out as previously described (15,20).

### Lipid overlay assays

PIP-Strips and Sphingo-Strips (Echelon biosciences) were used to test the lipid binding properties of GST-MAM^HS^. Membranes were blocked with blocking buffer (0.1% v/v Tween 20, 5% skim milk in PBS) for 1 hour at room temperature with gentle shaking. 10 µM of protein were added and incubated for 1 hour with shaking. Strips were washed with washing buffer (0.1% v/v Tween 20 in PBS) three times for 10 min each. GST antibody (1:1000 in blocking buffer) was applied and incubated for 1 hour at room temperature with shaking. Membranes were washed with washing buffer for 10 minutes; this step was repeated three times. The secondary anti mouse IgG-HRP antibody (1:5000 in blocking buffer) was added and incubated for 1 hour at room temperature with shaking. Strips were then washed with washing buffer three times for 10 min each and developed using Clarity ECL^TM^ Western substrate.

### Protein-protein and protein-lipid interaction plate assays

Protein binding to immobilized lipids or mucins was measured by a modified version of an indirect quantitative ELISA as follows: Mucin from porcine stomach type II, mucin from porcine stomach type III and mucin from bovine submaxillary glands (Sigma) were used to study the binding between MAM^HS^ and mucin. Mucin was dissolved in PBS at 4 °C for 20 hours with gentle shaking. For mucin immobolization, 100 µl of mucin at a concentration of 50 µg/ml was added into 96-well high-binding microtiter plates, plates were sealed and incubated at 4 °C for 20 hours. Plates were washed once with PBS, and 150 µl/well of blocking buffer (1% BSA in PBS) was added, plates were sealed and incubated at room temperature for 1 hour with gentle shacking. Then the wells were washed three times with PBS. Next, 100 µl/well of serial dilutions of MAM^HS^ in PBS (concentratinons as indicated in Figures) were added, plates were sealed and incubated for 2 hrs at room temperature with gentle shaking. The wells were washed three times with washing buffer. 100 µl/well of GST-antibody (1:1000 in blocking buffer) were added, plates were sealed and incubated for 1 hour at room temperature with gentle shaking. Wells were washed three times with washing buffer, and 100 µl /well of secondary anti-mouse IgG-HRP antibody (1:5000 in blocking buffer) were added, plates sealed and incubated for 1 hour at room temperature with gentle shaking. The wells were washed 3 times and 100 µl/well of Clarity ECL^TM^ Western substrate was added for detection. Bioluminescence was visualized on a BioRad Imaging system and quantified in a FLUOStar Omega plate reader. For experiments with desulfated mucin, mucin from porcin type II was desulfated with 100 µl of sulfatase from *Helix promatia* (Sigma) which was added at concentrations ranging from 0.1-3 U/ml in 20 mM sodium posphate buffer pH 5 and incubated at room temperature for 24 hours with gentle shaking. Following desulfation, mucin was washed 3 times with washing buffer and used for plate assays as described above. Alternatively, mucin was desulfated by incubation with cultures of *B. theta* wt *and B.theta* anSME mutant at OD 0.6 under anaerobic conditions at 37 °C for the indicated time points. Following desulfation, mucin was washed, blocked, and used for plate assays as described above.

To test binding of purified proteins to lipids, 3-O-sulfo-D-galactosyl-ß-1-1’-N-lignoceroyl-D-erythro-sphingosine, N-lignoceroyl-D-erythro-sphingosine and 1,2-dioleoyl-*sn*-glycero-3- phosphate (Avanti Polar Lipids) were dissolved in chloroform:methanol:water at a ratio of 2:1:0.1. 50 µl of lipids at a concentration of 200 µg/ml were immobilized in 96-well glass microtiter plates and left at room temperature for 20 hours for solvent evaporation. Control wells contained only solvents but no lipid. 150 µl/well of blocking buffer (1% BSA in PBS) was added, sealed and incubated at room temp for 1 hour. Wells were washed three times and protein binding was probed as described above for protein-protein interaction plate assays.

### Measurement of sulfate release from mucin

Mucins at a concentration of 50 µg/ml were desulfated as described above. In triplicate, 100 µl of supernatant from each reaction was transferred into a 96-well microtiter plate and 100 µl of a 20% barium chloride solution was added to each well. The optical density (OD_600_) was measured and converted into sulfate concentration using a standard curve of Na_2_SO_4_.

### Measurement of sulfatase activity

*B. theta* wt and *B. theta* anSME mutant were cultured in BHI containing 2.5 mg/ml mucin from porcine stomach type II and incubated anaerobically at 37 °C for 48 hours. The supernatant was collected after centrifuging cultures at 13000xg for 5 min. Serial dilutions from the supernatant were prepared using 0.2% NaCl. 100 µL of supernatant were incubated with 500 µL of 200 mM sodium acetate buffer and 400 µL of 6.25 mM p-nitrocatechol sulfate solution at 37 °C for 30 min. Then 5 ml 1 N NaOH was added to stop the reaction. The absorbance at 515 nm was detected using a Jenway 6300 UV/Vis spectrophotometer. Sulfatase from *Helix promatia* (Sigma) was used to prepare the standard curve.

### Protein pull-down experiments and protein identification

Hela cells (ATCC clone CCL-2) in confluent growth were harvested using a cell scraper and centrifuged at 1000xg, 22 °C for 5 minutes. The pellet was washed in ice cold buffer (20 mM Hepes-KOH pH 7.3, 110 mM KAc, 2 mM MgAc, 1 mM EGTA, 2 mM DTT), and 1ml of lysis buffer (10 mM Hepes-KOH pH 7.3, 10 mM KAc, 2 mM MgAc, 2 mM DTT, Roche protease cocktail) was added and left on ice for 10 minutes. Cells lysate was centrifuged at 12000xg for 12 min. 200 µg GST-MAM^HS^ or GST (control) was added to the cell lysate and incubated for 5 min at room temperature. DSP cross-linker in DMSO was added to give a final concentration of 100 µg/ml and incubated at room temperature for 30 min. Tris-HCl pH 8.0 was added to 10 mM to quench the reaction and the solution was added to glutathione sepharose beads, incubated for 2 hours at room temperature and then 20 hours at 4 °C. The suspension was centrifuged and the pellet was washed 4 times with washing buffer (20 mM Hepes-KOH pH 7.3, 110 mM KAc, 2 mM MgAc, 1 mM EGTA, 2 mM DTT, 0.1% Tween-20, 150-500 mM NaCl). Proteins were eluted into SDS sample loading buffer, boiled and resolved by SDS-PAGE. Gels were Coomassie stained and bands were cut out and sent for protein tryptic digest and identification of peptides by LC MS/MS on a Thermo Orbitrap Elite system coupled to a Dionex nano LC (University of Birmingham Advanced Mass Spectrometry Core).

### Bacterial attachment assays

Hela cells at a concentration of 1.5×10^5^ cells/well were seeded on coverslips 3 days before the experiments. Colorless DMEM containing *E.coli* BL21 expressing MAM^HS^ at an MOI of 100 was added to cells. For competition experiments, infection medium also contained 50 µg/ml mucin or desulfated mucin, or lactose-3-sulfate, lactose, N-acetyl glucose amine-6-sulfate, N-acetylglucosamine, galactose-6-sulfate or galactose at a concentration of 200 µM and was pre-incubated with bacteria for 1 hour prior to addition to cultured cells. Following bacterial adhesion, DMEM medium was removed and cells were washed three times with PBS to remove non-adherent bacterial cells, and 0.5% Triton 100 –X in PBS was added to lyse mammalian cells. Serial dilutions were prepared and cultured on LB agar at 37 °C for 24 hours to determine colony forming units. To visualize bacterial attachment to host cells, bacteria were co-transformed with pDP151 (mCherry) to give constitutive red fluorescence. DMEM medium was removed following the adhesion experiment and cells were washed three times with sterile PBS. For fixation, 3.7% paraformaldehyde in PBS was added for 15 min at room temperature. Cells were washed, permeabilized with 0.1% Triton X-100 in PBS for 5 min and stained with Hoechst and Alexa488- phalloidin for 10 min. Coverslips were mounted using Prolong Antifade mountant and imaged on a Zeiss Axio Observer Z1 microscope fitted with 40x/1.4 Plan Apochromat objective and ORCA-Flash4.0 camera. Images were acquired using ZEN 2.0.0.10 software and processed using ImageJ and Corel Draw X8 Graphics Suite.

### Mucin transmigration assays

Transwell filters (24-well thincert, 3.0µm pore diameter, Greiner Bio-One) were coated with 50 µL of 10 mg/ml mucin from porcine stomach type II at 4 °C for 20 hours, then placed onto 24-well plates containing 600 µl DMEM without phenol red. 100 µl of 10^6^ CFU/ml *E. coli* was added to the top well and incubated at 37 °C for 2 hours. Bacterial concentrations in the bottom well were enumerated by dilution plating on LB agar following incubation at 37 °C for 20 hours. To study the effect of sulfatase and *B. theta* on bacterial transmigration, *B. theta* wt and *B. theta* anSME mutant were grown anaerobically in BHI at 37°C to an OD_600_ of 0.6. Mucin was treated with 100 µl of culture supernatant from *B. theta* wt or *B. theta* anSME mutant at 37°C for 24 hours. Alternatively, mucin was treated with *Helix promatia* sulfatase as described above, prior to use in transwell assays.

## ACKNOWLEDGMENTS

We would like to thank Elisabeth Lowe (Newcastle University) and Eric Martens (University of Michigan) for providing *B. theta* strains, our collaborator Meera Unnikrishnan (University of Warwick) for providing HT-29 MTX cells, Joao Correia (University of Birmingham) for technical assistance with fluorescence microscopy, Jinglei Yu (University of Birmingham) for technical assistance with mass spectrometry, and members of the Krachler lab for critical reading and comments on the manuscript. A.M.K., D.V. and D.H.S. were funded through BBSRC grants BB/L007916/1 and BB/M021513/1. F.A. was funded through an Iraq Ministry of Higher Education and Scientific Research Postgraduate Scholarship.

## AUTHOR CONTRIBUTIONS

F.A, D.P.V, D.H.S and A.M.K did experiments and analyzed data. F.A and A.M.K drafted the manuscript. All authors read, edited and agreed to the final version of the manuscript.

